# Curiosity-Motivated Incidental Learning With And Without Incentives: Early Consolidation And Midbrain-Hippocampal Resting-State Functional Connectivity

**DOI:** 10.1101/2022.12.23.521819

**Authors:** Stefanie Meliss, Kou Murayama

**Author notes:** Correspondence should be addressed to Kou Murayama, Hector Research Institute of Education Sciences and Psychology, University of Tübingen, Tübingen, 72072 Germany, and Stefanie Meliss, School of Psychology and Clinical Language Sciences, University of Reading, Reading, Berkshire, RG6 6AH, UK,. **Author Note**. Scripts used to process and analyse the data were uploaded to GitHub (https://github.com/stefaniemeliss/MIC_paper). The repository further contains the behavioural data. Raw and pre-processed fMRI were uploaded on OpenNeuro (https://doi.org/10.18112/openneuro.ds004182.v1.0.0). ROI masks and unthresholded fMRI results at group level are available as a NeuroVault collection (https://neurovault.org/collections/13320/).

## Abstract

Human memory is selective and not all experiences are remembered. Both monetary rewards/incentives and curiosity have been found to motivate and facilitate learning by dopaminergic midbrain projections to the hippocampus during encoding. In this study, we examined potential brain mechanisms during early consolidation period that jointly or independently contribute to these facilitating effects. Participants (N = 50) watched 36 videos of magic tricks and rated their “subjective feelings of curiosity” while the availability of extrinsic incentives was manipulated between groups. Functional magnetic resonance imaging (fMRI) data were acquired before, during, and after learning, and memory for magic tricks was assessed one week after. Our analysis focused on the change in resting-state functional connectivity (RSFC) between the dopaminergic midbrain and the anterior hippocampus, a dopaminergic consolidation mechanism previously reported in the context of extrinsically motivated learning. Changes in RSFC were correlated with behavioural measures of learning, i.e., the total number of items encoded and the curiosity-driven memory benefit. We found that brain-behaviour correlations differed depending on the availability of extrinsic incentives. More specifically, the correlation between the total number of items encoded and RSFC change was significantly different in the incentivised compared to the control group. The curiosity-driven memory benefit, however, did not correlate with changes in RSFC in either of the groups. In sum, this suggests that curiosity-motivated learning might be supported by different consolidation mechanisms compared to extrinsically motivated learning and that extrinsic incentives influence consolidation mechanisms supporting learning.

**Key points:**

- A new curiosity-motivated incidental encoding paradigm was used to investigate how dopaminergic consolidation mechanisms support learning and whether this is further influenced by the availability of monetary incentives.
- Changes in resting-state functional connectivity between the dopaminergic midbrain and the anterior hippocampus, a dopaminergic consolidation mechanism, predicted learning outcomes significantly differently if monetary incentives were available.
- These results might suggest that learning motivated by curiosity might rely on different neural mechanisms during early consolidation than learning motivated by monetary incentives.

---

Consolidation describes processes during which a newly formed memory trace becomes increasingly stabilised and thereby transformed into long-term memory (Dudai, 2012; Wang & Morris, 2010). Both, cellular-level consolidation (i.e., selective modification of synapses within the hippocampus (HPC)) as well as subsequent systems-level consolidation (i.e., stabilisation of memory traces through cross-regional, HPC-cortico interactions) are supported by replay and/or reactivation processes (recently reviewed by Tambini & Davachi, 2019) where sharp wave ripples originating in the HPC facilitate the coordinated reactivation of cortical traces through oscillations (Cowan et al., 2021). Reactivation is likely related to the lingering excitability of the neurons that were engaged during encoding and synaptic plasticity processes (for a detailed explanation, see Atherton et al., 2015). Importantly, it has been shown that reactivation manifests in changes in the blood-oxygen-level-dependent (BOLD) signal and in functional connectivity (FC) patterns over minute-long timescales (Logothetis et al., 2012; Tambini & Davachi, 2019). This allows brain mechanisms related to early consolidation to be studied in MRI paradigms using a baseline (or pre-learning) scan, a memory task, followed by a memory task and a post-learning scan (Deuker et al., 2013; Gruber et al., 2016; Kukolja et al., 2016; Murty et al., 2017, 2019; Tambini et al., 2010; Tambini & Davachi, 2013). Changes from pre- to post-learning (e.g., increased FC) can be interpreted as related to the ongoing strengthening of memory traces, especially when related to encoding (Tambini & Davachi, 2019). Research focusing on changes in RSFC between the HPC and other brain areas from pre- to post-learning rest and how they relate to learning provide further support for their role in memory (Murty et al., 2017; Schlichting & Preston, 2014; Tompary et al., 2015).

## Monetary Incentives and Memory Consolidation

The neurotransmitter dopamine seems to exert effects on the memory consolidation processes by influencing late long-term potentiation (LTP) processes (Lisman et al., 2011) which has been discussed as the mechanism of motivated learning (Miendlarzewska et al., 2016; Shohamy & Adcock, 2010). Behavioural studies often report enhancing effects of monetary incentives on encoding only in delayed, but not immediate memory tests (Murayama & Kitagami, 2014; Murayama & Kuhbandner, 2011; Patil et al., 2017; Wittmann et al., 2005). In animal research, more sharp wave ripples were recorded in the rodent HPC following reward-motivated learning compared to the absence of reward, implying that the reactivation patterns in the HPC might reflect experiences associated with reward (Singer & Frank, 2009). Intriguingly, reactivation has also been observed in recordings in the ventral tegmental area (VTA) during awake rest and sleep after receiving different rewards (Valdés et al., 2015). In addition, following appetitive spatial learning, replay in the HPC coincided with reward-responsive neurons in the VTA during an awake rest, but not during sleep (Gomperts et al., 2015). If dopaminergic projections to the HPC are stimulated during the exploring of novel environments or encoding spatial encoding using optogenetics, this is associated with enhanced reactivation of pyramidal cells in the HPC and better recall indicating that activity in dopaminergic cells can enhance hippocampal replay (McNamara et al., 2014).

Human fMRI studies also suggest a dopaminergic influence on memory consolidation in reward learning in HPC, VTA/SN and their interaction. Gruber and colleagues (2016) presented a reward-manipulated incidental item-associative encoding task to participants inside the MRI and also collected fMRI data during pre- and post-learning rest. Results of the immediate memory test indicated that reward enhanced encoding of associative information. The authors showed that in the absence of an overall increase in RSFC between HPC and midbrain from pre- to post-learning rest, individual differences therein positively predicted the reward-driven associative memory effect. That interindividual differences in changes in RSFC between HPC and midbrain can predict later associative memory for information presented in high-reward contexts was recently replicated in an intentional encoding paradigm with a memory test after 24h (Cohen et al., 2021). In sum, the results provide evidence that monetary rewards influence neural post-encoding processes supporting the preferred consolidation of stimuli associated with rewards.

Trying to understand the mechanisms for the observed effects of changes in HPC-VTA RSFC on encoding, Murty and colleagues (2017) used a incentive-motivated learning paradigm with word-image pairs in which high or low value was associated with the category of the image. The fMRI analysis focused on changes in RSFC from pre- to post-learning between anterior HPC (aHPC) or VTA and the category-selective cortex used in the high-reward condition. The authors showed that changes in RSFC changes between category-selective cortex and aHPC or VTA, respectively, positively predicted high-value associative memory, but had no effect on low-value associative memory, thereby suggesting that memory for high-value information is supported by mechanisms that specifically target the sensory cortex in which the information was initially processed, potentially via neuromodulation of relevant cortices that might have received a behavioural tag during encoding (Moncada et al., 2015; Murty et al., 2017). Taken together, the literature reviewed here indicates that dopamine supports adaptive memory for events with motivational salience not only during encoding, but also during consolidation (Cowan et al., 2021; Shohamy & Adcock, 2010).

## Curiosity and Memory Consolidation

Recent studies have also shown that the same midbrain neurons that respond to the anticipation of primary extrinsic rewards also respond to the anticipation of non-instrumental information (Bromberg-Martin & Hikosaka, 2009) – a concept closely related to curiosity (Gottlieb & Oudeyer, 2018). These studies suggest that information is an intrinsic reward (FitzGibbon et al., 2020; Marvin & Shohamy, 2016) that can modify behaviour in a similar vein as extrinsic rewards. In fact, the literature seems to demonstrate a general overlap between the effects of curiosity and extrinsic rewards/incentives on encoding as (1) both are associated with the release of dopamine, (2) both show behavioural memory enhancement effects, that are (3) both supported by activity in the HPC and the VTA/SN as well as the interaction between them during the encoding of new information. This conjecture is also supported by the evidence that suggests that the effects of novelty on memory are supported by consolidation-related processes (Ballarini et al., 2013; Fenker et al., 2008; Lisman & Grace, 2005; Moncada & Viola, 2007; Wang et al., 2010), and novelty has been considered as a critical factor influencing curiosity (Berlyne, 1950; Ryan & Deci, 2000).

However, the evidence also suggests potential differences between curiosity- and extrinsically-motivated learning. For example, behavioural evidence showed that the effects manifest on different time scales. More specifically, incentive/reward effects on encoding are frequently found after long, but not short, delays between encoding and retrieval as evidenced by studies comparing the effects in immediate and delayed memory tests (Murayama & Kitagami, 2014; Murayama & Kuhbandner, 2011; Patil et al., 2017; Wittmann et al., 2005). This is in line with proposals that the effects of monetary reward on encoding are dopaminergic, influencing late LTP and are hence only apparent in delayed, but not immediate encoding (cf. Gruber et al., 2016; Murty & Adcock, 2014). The effects of curiosity on encoding, on the other hand, have consistently been found after short (Brod & Breitwieser, 2019; Galli et al., 2018; Gruber et al., 2014; Jepma et al., 2012; Ligneul et al., 2018; Mullaney et al., 2014; Murphy, Dehmelt, et al., 2021; Poh et al., 2021) and long (Duan et al., 2020; Fastrich et al., 2018; Gruber et al., 2014; Halamish et al., 2019; Kang et al., 2009; Marvin & Shohamy, 2016; Murayama & Kuhbandner, 2011; Swirsky et al., 2021) delays between encoding and retrieval. A study examining the effects of curiosity levels on later memory by testing half of the items after a short and the other half of the items after a long delay found that curiosity correlated with encoding at both time intervals (McGillivray et al., 2015). Likewise, another study did not observe any effects of delay on curiosity-motivated learning (Stare et al., 2018).These findings were somewhat puzzling because if the effect of curiosity on encoding was purely/predominantly dopaminergic as it is assumed to be the case for monetary rewards (Miendlarzewska et al., 2016; Shohamy & Adcock, 2010), cellular models of learning and the effects of dopamine and late LTP (Lisman et al., 2011) would predict that the effects of curiosity on memory should only be seen after long delays (allowing for the expression of late LTP) or that they would at least be more pronounced after long compared to short delays. Hence, the presence of memory-facilitating effects of curiosity after short and long delays indicates that reward- and curiosity-motivated learning might be supported by differential neural mechanisms.

## Current Research

The current research aims to investigate whether curiosity-motivated learning is supported by similar post-encoding systems-level consolidation processes (i.e., change in RSFC between HPC and VTA/SN) as have been reported for extrinsically motivated learning. While post-encoding mechanisms of reward-motivated learning have been studied and related to the reactivation of local activation patterns within HPC and VTA/SN (Gomperts et al., 2015; Gruber et al., 2016; McNamara et al., 2014; Murty et al., 2017; Singer & Frank, 2009; Valdés et al., 2015), to the best of our knowledge, research on post-encoding mechanisms of curiosity-motivated learning is thus far lacking. Due to the involvement of overlapping brain regions during encoding, it is tempting to assume that post-encoding processes would also be overlapping. However, the different temporal contingencies of curiosity and incentive/reward effects allow for the possibility that different consolidation processes support these effects.

Given the role of dopamine, VTA/SN, and HPC during curiosity-motivated encoding as well as its conceptual partial overlap with monetary incentives/rewards and novelty, we investigated whether similar post-encoding mechanisms as previously reported in the context of reward and novelty can also be found for curiosity and were interested in post-encoding systems-level interactions between HPC and VTA/SN. More specifically, we focused on the aHPC because the aHPC has previously been implicated in post-encoding processes of high-value information (Murty et al., 2017). To elicit curiosity, magic trick video stimuli from a validated stimulus database (Ozono et al., 2021) were used that participants incidentally encoded during an incentivised orientation task inside the MRI scanner, accompanied by pre- and post-learning resting-state scans. The availability of monetary incentives was manipulated in a between-subject design, in order to calculate the post-encoding FC change separately when rewards were available and when not. Our research questions were the following: (1) Does the RSFC between VTA/SN and aHPC increase as a function of learning? (2) Does the availability of extrinsic incentives influence the change in RSFC between VTA/SN and aHPC as a function of learning? (3) Do individual differences in the change in RSFC between VTA/SN and aHPC predict behavioural measures of learning (i.e., the totaal number of items encoded and the curiosity-driven memory benefit) and is this association sensitive to the availability of extrinsic incentives?

## Materials & Methods

The MRI dataset analysed here has been made publicly available as Magic, Memory, and Curiosity (MMC) Dataset (https://doi.org/10.18112/openneuro.ds004182.v1.0.0) and the present work focuses on the pre- and post-learning fMRI data (for an analysis of the task data, please see Meliss, van Reekum, et al., 2022). The methods of data collection and processing are summarised here (for a more detailed account, please refer to the dataset descriptor; Meliss, Pascua, et al., 2022).

### Participants and Design

The sample included data from 50 healthy adults (36 (or 72%) female) aged 18-37 (*M* = 25.32, *SD* = 5.19), randomly assigned to control and incentives groups. Participants were recruited using leaflets distributed across campus and leisure centres. All participants had normal or corrected-to-normal vision using contact lenses, were right-handed, fluent in English, did not suffer from any chronic illness, psychiatric disorders or cognitive impairments and did not take any psychoactive drugs. Women were only included if they were not pregnant or nursing.

The data collection consisted of three sessions: a pre-scanning session conducted online (not described here), an incentive- and/or curiosity-motivated incidental encoding and consolidation session inside the MRI scanner, and a surprise memory test online approximately a week later. The study used a between-subject design where the availability of extrinsic, monetary incentives for the performance in an incentivised orientation task was manipulated: in the incentives group, participants were instructed that each correct answer in the orientation task was worth an additional, performance-dependent bonus of £ 0.80 for each correct answer per trial (see below). Such instructions were not provided to participants in the control group. Importantly, performance feedback was not provided in either group.

Participants were compensated £ 30 for their participation and received an additional bonus payment of £ 7.20 (chance level performance in four-alternative forced choice orientation task across 36 trials), regardless of the group they are assigned to and regardless of their performance in the orientation task. The study design was approved by the University Research Ethics Committee (UREC) of the University of Reading (UREC 18/18).

### Material

To induce curiosity inside the MRI scanner, short videos of magic tricks from a publicly available stimulus collection (“Magic Curiosity Arousing Tricks (MagicCATs)”; Ozono et al., 2021) were used. These stimuli reliably induce curiosity and other epistemic emotions with sufficient within-person variability. For the purpose of this study, 36 magic tricks were selected as stimuli to be presented in the incidental incentive- and/or curiosity-motivated learning task based on the following criteria: (1) duration between 20 and 60 s, (2) various ranges of distinguishable materials and features, (3) broad range of average curiosity ratings as reported in the stimulus collection. The final selection of stimuli was edited using Adobe® Premiere Pro CC® (2015) software. The aim was to hide the faces of the four magicians performing the tricks as much as possible and to ensure similar viewing foci and dark backgrounds across videos. Where needed, additional editing was performed, e.g., removing subtitles. Importantly, all videos were muted, but still understandable due to the non-verbal nature of magic tricks. An individual mock video (duration 6 s) was created and added to the beginning of each magic trick where the first frame of the trick was displayed, overlaid with a black video. The viewing focus of that black video slowly opened to match the viewing focus of the video. The final magic trick files (1280×720 pixels) were on average 38.5 s long (*SD* = 8.63, min = 26.6, max = 58.64).

For the surprise cued memory test, a frame from each magic trick was extracted as a cue image (1920×1080 pixels). A frame was chosen, ensuring that it was prior to the moment of expectancy violation/surprise to not reveal the trick entirely, yet distinct enough to cue memory of the magic trick.

### Task Procedures

The paradigm consisted of different phases (see Figure 1): (1) a pre-learning rest phase (i.e., baseline), (2) an incentive- and/or curiosity-motivated incidental learning phase, (3) a post-learning rest phase (i.e., early consolidation), and (4) a delayed, surprise memory test. While phases 1-3 happened consecutively during the same session inside the MRI scanner (using Psychophysics Toolbox; Brainard, 1997 implemented in Matlab (2018b)), the fourth phase took place approximately a week later (*M* = 7 d 10 h 19 min, *SD* = 13 h 41 min) in a separate session.

**Figure 1.**
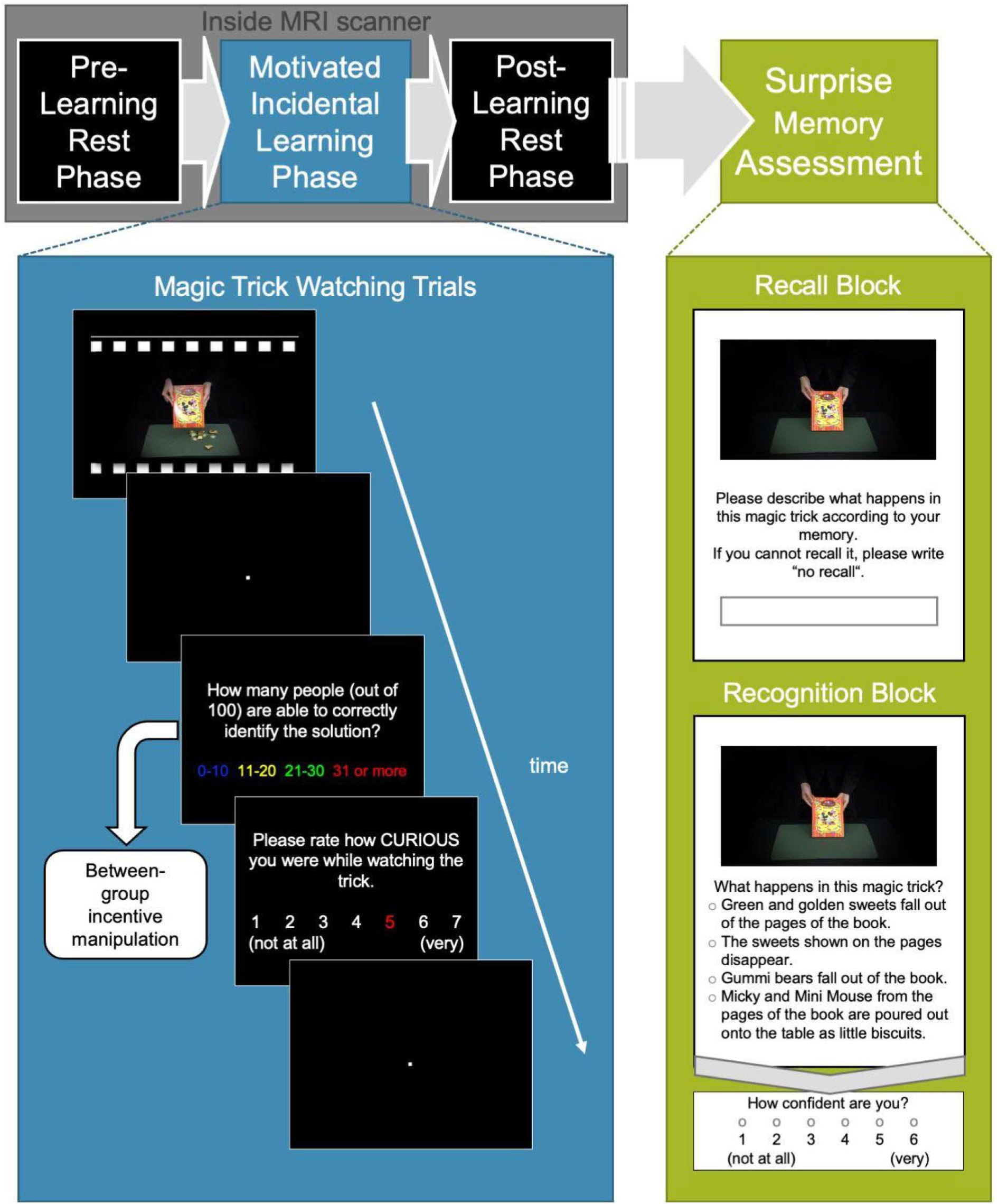
Task Procedures *Note*. The figure illustrates the structure of the different phases. Inside the MRI scanner, resting-state data were collected before the incidental encoding task (trial structure shown in the blue box on the left). This was followed by another resting-state scan. A week later, memory was accessed in a surprise test online (trial structure illustrated in the green box on the right).

#### Pre- and Post-Learning Rest Phases

During the rest phases (10 min each), participants were lying inside the MRI scanner and were monitored with an eye-tracking camera. A white rectangle spanning 90% of the height and width of the black screen was presented. Participants were asked to try and keep as still as possible while simply looking at the white screen, allowing blinking as usual. They were instructed that their brain activity at rest was measured and that they should hence try to not think about anything at all.

#### Motivated Incidental Learning Phase

During the incidental learning phase, participants viewed in total 36 magic tricks in pseudo-randomised order distributed across three blocks. Self-paced breaks were offered in between each block. The magic tricks and subsequent ratings were presented on a black background. The display of each magic trick was aligned with the scanner’s TTL (transistor-transistor logic) pulse. Each magic trick presentation was followed by a fixation (jittered between 4 and 10s; start TTL pulse aligned). Then participants were asked to give an estimate of how many people (out of 100) would be able to correctly figure out the solution to the magic trick and were presented with four options (‘0-10’, ‘11-20’, ‘21-30’, and ‘31 and more’) each one corresponding to a button on the four-button MRI-compatible response device and the printed in the corresponding colour. Importantly, participants in the incentives group were instructed before the start of each task block (but after the pre-learning rest phase) that each correct estimate would translate into an additional bonus of £ 0.80. This instruction was not presented to the control group and did not affect the trial structure itself which was the same in both groups. Participants’ response window was fixed to a duration of 6s. If a response was given before the end of the response window, all coloured writing was removed from the screen. After a fixed fixation (0.05 s), participants were asked to rate how curious they were while watching the magic trick on a scale from 1 (‘not curious at all’) to 7 (‘very curious’) by moving a randomly highlighted number on the curiosity scale to the left or right until it represented the chosen rating. This was then confirmed via a button press. The fixed response window here was 5.95 s and the font would again turn white if the rating was collected before the end of the response window. After another fixation was presented (jittered between 4 and 10 s), the next trial began.

#### Surprise Memory Assessment

During recruitment, participants were informed that there would be a follow-up assessment, taking place online (implemented using a developmental version of Collector; Haffey et al., 2020) a week after scanning. However, the purpose (i.e., memory test) was not communicated in advance. When asked about their guess regarding the hypothesis behind the study at the end of the MRI session, no participant mentioned learning, memory, or encoding. Reminders of the assessment were sent out two days in advance and participants received the link for the surprise memory assessment at the same time as their initial slot. The surprise memory test consisted of a recall and recognition block during which the same cue images were displayed in random order. In the cued recall block (data not reported here), participants were asked to describe what had happened in this magic trick according to their memory but were also instructed to enter ‘no recall’ if they could not remember a trick. During the cued recognition, they were presented with four descriptions of what could have happened in the magic trick and asked to select one and afterwards rate their confidence in the answer on a scale from 1 (‘not confident at all’) to 6 (‘very confident’).

### Behavioural Analysis

Behavioural analyses were conducted in R version 3.6.3 (R Core Team, 2020). The recognition data were dummy-coded by comparing the chosen answer against the correct answer, creating a recognition measurement regardless of confidence to allow us to test whether the performance in the recognition test was significantly above chance level (i.e., 25% given the four answer alternatives). Recognition performance was further combined with the confidence ratings (i.e., correct answer chosen with a confidence larger than 3) creating a high confidence recognition measurement reflecting recollection-based recognition memory (Yonelinas, 2001b, 2002) that was used as the main measurement of encoding in our analyses. This threshold was chosen as it showed the strongest between-group effect on encoding in behavioural pilot studies (Meliss & Murayama, 2019).

To relate FC measurements (see below) to behavioural measures of learning, two indices per subject were computed: the total number of items encoded and the curiosity-driven memory benefit. The total number of items encoded (i.e., recognised with confidence ratings above 3) was determined by calculating the sum of magic tricks encoded. To quantify the effects of curiosity on memory, we adopted and modified calculations of the ‘*curiosity-driven memory benefit*’ proposed by Gruber and colleagues (2014). More specifically, for each subject, the mean curiosity rating across all 36 trials was calculated and each trial was categorised depending on whether the respective curiosity rating was above or below the mean (of the participant) to create high- and low-curiosity trials. We then computed (1) the ratio between the number of magic tricks encoded eliciting high ratings of curiosity and the total number of magic tricks eliciting high ratings of curiosity; and (2) the ratio between the number of magic tricks encoded eliciting low ratings of curiosity and the total number of magic tricks eliciting low ratings of curiosity; to then (3) calculate the difference between these values. As such, positive values indicate that more magic tricks eliciting high than low ratings of curiosity were encoded and vice versa for negative values.

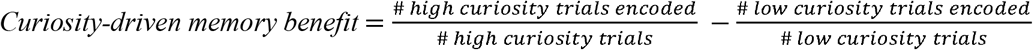

### fMRI Acquisition and Pre-processing

A 3.0T Siemens Magnetom Prisma scanner with a 32-channel head coil was used to acquire anatomical and functional images as well as a B_0_ field map. To restrict excessive head motion, the participants’ head was padded inside the head coil. An Echo-Planar Imaging (EPI) sequence was used to obtain whole-brain (37 axial slices, 3 × 3 × 3 mm, interslice gap of 0.75mm) T2*-weighted images (repetition time (TR) = 2000ms, echo time (TE) = 30ms, field of view (FoV): 1,344 × 1,344 mm2, flip angle (FA): 90°; phase encoding direction: P>>A). Additionally, B_0_ data were acquired immediately after the pre-learning rest on the same image matrix and the same geometric prescription as the functional data, using a dual-TE 2D gradient-echo sequence (TR = 488 ms, TE1/TE2 = 5.19/7.65 ms, FA = 60°). Lastly, a high-resolution T1-weighted whole-brain image (192 × 1-mm slices) was acquired using an MPRAGE-gradient sequence (in-plane resolution of 1 × 1 × 1 mm; TE: 2.29 ms; TR: 2300 ms; inversion time (TI): 900 ms; FOV: 240 × 240; FA: 8°). Instructions and stimuli were displayed via back projection using a mirror attached to the head coil above the eyes. While inside the MRI scanner, an MRI-compatible eye tracker was used to allow the experimenter to monitor whether the participants kept their eyes open during the rest phases and whether they attended the stimuli during the motivated incidental learning phase.

Imaging data were pre-processed using AFNI (version 21.2.03; Cox, 1996) and the same pipeline was applied to data collected during rest and encoding. Steps included B_0_ distortion correction, despiking, slice-timing and head motion correction and normalisation to MNI space using the ICBM 2009c Nonlinear Asymmetric Template. Pre-processed data were smoothed to achieve an approximate, uniform smoothness of full width half maximum (FWHM) kernel of 4 mm using AFNI’s *‘3dBlurToFWHM’*. Compared to conventional smoothing where a Gaussian kernel with a specific FWHM is applied, 3dBlurToFWHM iteratively smooths the EPI time series until the images have reached the desired uniform smoothness within the specified mask (Scheinost et al., 2014).

Following smoothing, time series of each voxel were scaled to a mean of 100. Local white matter time series, the first three principal components of the lateral ventricles as well as motion estimates were included as regressors of no interest to denoise the data. During linear regression, time courses were also band-pass filtered for frequencies between 0.01 and 0.1 Hz. Time points were censored (i.e., set to zero) if the Euclidean norm of per-slice motion exceeded 0.3mm or if more than 10% of brain voxels were outliers.

The time series acquired during the rest phases consisted of 300 volumes each. For each of them, the global correlation (GCOR; Saad et al., 2013) was computed using AFNI’s *‘@compute_gcor’*. GCOR represents a single value for each dataset, created by computing the average correlation over all possible combinations of voxels within the grey matter mask, and then averaging all values within the mask. As such, GCOR captures brain-wide correlations and between-subject fluctuations therein are assumed to be driven by sources of noise.

### Method to Define Regions-of-Interest (ROI) Masks

Due to the established roles of HPC and VTA/SN as well as their connectivity in motivated learning in the context of incentives and curiosity have previously been established (Adcock et al., 2006; Cohen et al., 2021; Gruber et al., 2014, 2016; Kahn & Shohamy, 2013; Murphy, Ranganath, et al., 2021; Murty & Adcock, 2014; Poh et al., 2021; Wolosin et al., 2012), we aimed to determine how their FC during post-encoding rest predicts behavioural measures of learning. Because evidence suggests that activity during encoding and consolidation in support of remembering is predominantly centred in anterior parts of the HPC (Kim, 2011; Murty et al., 2017; Spaniol et al., 2009), the analysis focused on the aHPC and the same masks were used as described in previous work (Meliss, van Reekum, et al., 2022). The bilateral HPC mask was extracted from the Glasser Human Connectome Project atlas (Glasser et al., 2016) and the aHPC was determined based on the uncal apex using MNI coordinate y = 21P (Poppenk et al., 2013). The mask for the VTA/SN was created using *atlaskit* (https://github.com/jmtyszka/atlaskit) by (1) extracting SN pars reticulata (SNr), SN pars compacta (SNc), and VTA from a high-resolution probabilistic subcortical nuclei atlas in MNI space (Pauli et al., 2018) specifying a probability threshold of 15% and then (2) combining all three into a single mask. The masks were shown in Figure 2 and can be accessed in the NeuroVault collection (https://neurovault.org/collections/13320/).

**Figure 2.**
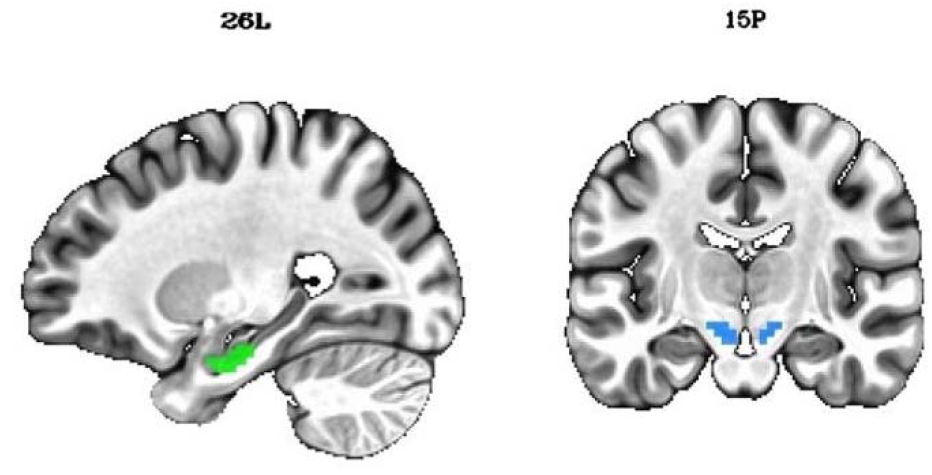
aHPC and VTA/SN Seeds *Note*. aHPC is shown in green, whereas VTA/SN is shown in blue.

### Functional Connectivity Analysis

Similar to Gruber and colleagues (2016), the main focus of this work was to determine how FC between the *a priori* defined seed regions aHPC and VTA/SN (i.e., RSFC) changes from pre- to post-learning. For each pre-processed rest time series separately, AFNI’s *‘3dSetupGroupInCorr’* was specified, loading in the respective time series of all subjects into an object to be used as input for *‘3dGroupInCorr’*, a program that in turn computes the average time series within the specified input mask (i.e., aHPC) to then compute the Pearson correlation between the averaged time course within the input mask and the time course of each voxel in the brain. GCOR was included as a subject-level covariate to further account for nuisance sources. *‘3dGroupInCorr’* does not only calculate seed-based FC at group level but also outputs the individual Fisher’s *z*-transformed results. Unthresholded group level results as well indiUnthresholded statistical maps were uploaded to NeuroVault (https://neurovault.org/collections/13320/). From the individual seed-based maps, the values of voxels within the VTA/SN mask were extracted and averaged as a measurement of RSFC between aHPC and VTA/SN at pre- and post-learning, respectively. Change in RSFC was computed by calculating the difference between both to be correlated with behavioural measures of learning (Gruber et al., 2016). This was done across the whole sample and within each group separately. The *corcor* package (Diedenhofen & Musch, 2015) was used to test differences in brain-behaviour-correlations between groups for significance.

### Exploratory Analysis

The main analysis focused on FC between the *a priori* defined seed regions aHPC and VTA/SN. However, whole-brain seed-based RSFC analyses were conducted for exploratory purposes. For this, data from pre- and post-learning rest phases from each subject were loaded into ‘3dGroupInCorr’, specifying aHPC and VTA/SN as seeds, respectively, to compute two seed-based FC maps, one for each ROI as seed, while controlling for GCOR to further account for sources of noise at group level, creating two seed-based maps per subject where the value in each voxel represents the Fisher’s z-transformed correlation coefficient with the averaged time series of each seed, respectively. Individual Fisher’s z-transformed maps were extracted for the pre- and post-learning resting phase to compute the RSFC as the difference between post- and pre-learning rest. These individual difference maps were then used as input for Two Sample t-tests to determine any differences between the groups with respect to changes in RSFC with either seed. Additionally, the difference maps were submitted to ANCOVA models, specifying the behavioural measures of learning as covariates in separate models to identify voxels showing a significant difference in slopes with respect to the covariate between the groups. To determine clusters of voxels exhibiting a significant change in RSFC with either of the seeds, a cluster-defining threshold of p = 0.001 and k = 5 were used (Woo et al., 2014). Unthresholded statistical maps were uploaded to NeuroVault (https://neurovault.org/collections/13320/).

## Results

### Behavioural Measures of Learning

To determine whether participants’ performance in the recognition memory test exceeded chance level (i.e., 25% of 36 items), the sum score in recognition performance regardless of confidence was calculated for each participant. A one-tailed t-test showed that the performance in the recognition test (*M* = 22.54, *SD* = 4.12, min = 15, max = 32) was significantly above chance level, *t*(49) = 23.26, *p* < 0.001, *d* = 3.29, 95% CI [2.42; 4.16] and no subject showed individual sum scores below chance level. Together, this suggests participants indeed incidentally encoded the magic tricks.

Encoding was measured as high confidence recognition, (i.e., recognition with a confidence rating > 3), which is a recollection-based recognition memory measurement (Yonelinas, 2001, 2002). Across both groups, the average total number of items encoded was 15.14 (*SD* = 5.35, min = 3, max = 27) and no significant differences (*M*_*C*_ = 15.52, *SD*_*C*_ = 4.65, *M*_*I*_ = 14.76, *SD*_*I*_ = 6.04) were observed, *t*(45.03) = 0.50, *p* = 0.621, *d* = 0.42, 95% CI [-0.15; 0.99].

To derive individual coefficients quantifying the curiosity-driven memory benefit (Gruber et al., 2014), comparing the number of items later remembered vs. forgotten in states of high vs. low curiosity. Across both groups, the average curiosity-driven memory benefit was 0.02 (*SD* = 0.22, min = -0.86, max = 0.36) and a significant difference between the groups was observed, *t*(47.91) = 2.139, *p* = 0.038, *d* = 0.61, 95% CI [0.02; 1.19], indicating higher curiosity-driven memory benefit scores in the control compared to the incentives group (*M*_*C*_ = 0.08, *SD*_*C*_ = 0.21, *M*_*I*_ = -0.05, *SD*_*I*_ = 0.22). This group effect remained significant after removing two outliers (as defined by the interquartile range method), *t*(43.03) = 2.488, *p* = 0.017, *d* = 0.72, 95% CI [0.19; 1.31].

### Resting-State Functional Connectivity (RSFC) Between aHPC and VTA/SN

RSFC between aHPC and VTA/SN was quantified separately for pre- and post-learning rest. While RSFC values increased slightly numerically from pre- (*M* = 0.035, *SD* = 0.025) to post-learning rest (*M* = 0.039, *SD* = 0.030), the change from pre- to post-learning (aHPC-VTA/SN-RSFC change; *M* = 0.004, *SD* = 0.031) was not significantly larger than zero, *t*(49) = 0.931, *p* = 0.178, *d* = 0.14, 95% CI [- 0,44; 0.70]. Likewise, while the difference was numerically larger in the control (*M* = 0.007, *SD* = 0.032) compared to the incentives group (*M* = 0.006, *SD* = 0.030), the group difference was also not significant, *t*(47.816) = 0.785, *p* = 0.435, *d* = 0.22, 95% CI = [-0.35; 0.79].

### Brain-Behaviour Correlations

The primary objective of our analysis was to determine whether changes in RSFC between aHPC and VTA/SN were associated with behavioural measures of learning and whether this was influenced by the availability of extrinsic incentives. We, therefore, correlated behavioural measures of learning (i.e., the total number of items encoded and curiosity-driven memory benefit) with aHPC-VTA/SN-RSFC change, across the whole sample and within each group separately. The patterns were illustrated in Figure 3.

**Figure 3.**
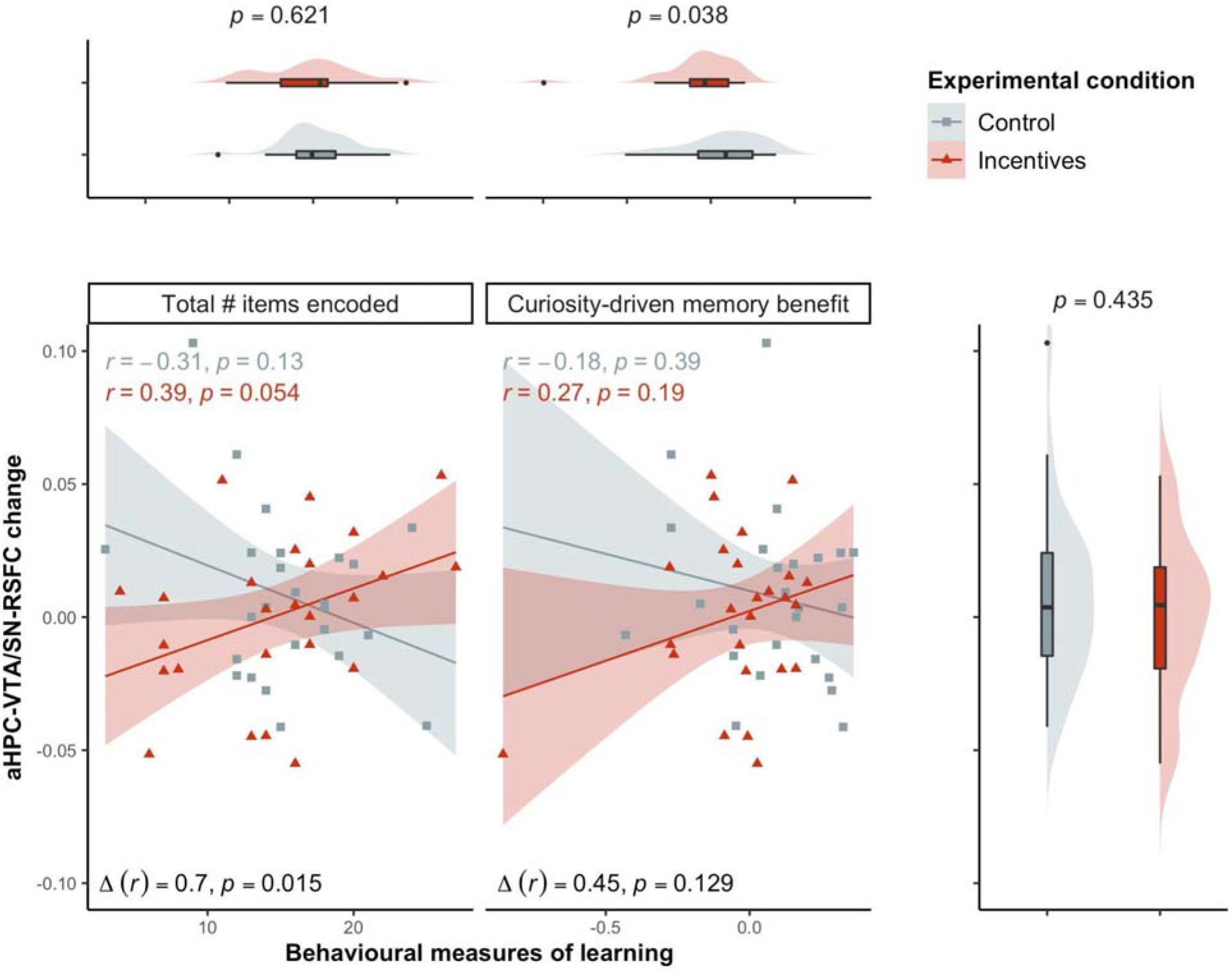
Correlation Between aHPC-VTA/SN-RSFC Change and Behavioural Measures of Learning *Note*. The scatter plots above show the association between behavioural measures of learning on the x-axis and changes in RSFC between aHPC and VTA/SN from pre- to post-learning rest on the y-axis. Data are shown separately for the total number of items encoded (left) and the curiosity-driven memory benefit (right). Above the scatter plots, distribution and box plots were added to further illustrate the distribution of the behavioural measures of learning in each group and likewise for the change in RSFC on the right. Please note that the data for the behavioural measures were scaled before creating distribution and boxplot to allow comparability. Different colours were used for the control and the incentives group, respectively. Regression lines with 95% confidence have been added to illustrate the relationship between the behavioural measures of learning and aHPC-VTA/SN-RSFC change within each group together with the corresponding correlation coefficients. Group effects in behavioural measures and RSFC change as well as in brain-behaviour-correlations were further added (without removing outliers).

While the Pearson’s correlation coefficient between the total number of items encoded and aHPC-VTA/SN-RSFC change was close to zero across the whole sample, *r* = 0.08, *t*(48) = 0.557, *p* = 0.581, 95% CI [-0.20; 0.35], the Pearson’s correlation coefficient was numerically positive within the incentives group, *r*_*I*_ = 0.39, *t*(23) = 2.028, *p* = 0.054, 95% CI [-0.01; 0.68], but numerically negative in the control group, *r*_*C*_ = -0.31, *t*(23) = -1.579, *p* = 0.128, 95% CI [-0.63; 0.09]. Importantly, the difference in correlation between both groups was significant, Δ*r*_*C<I*_ = 0.70, *z* = 2.436, *p* = 0.015, 95% CI [0.13; 1.13].

Similar patterns were observed in the correlation between curiosity-driven memory benefit and aHPC-VTA/SN-RSFC change. Again, the correlation coefficient was numerically positive within the incentives group, *r*_*C*_ = 0.27, *t*(23) = 1.34, *p* = 0.193, 95% CI [-0.14; 0.60], and numerically negative in the control group, *r*_*I*_ = -0.18, *t*(23) = -0.878, *p* = 0.388, 95% CI [-0.53; 0.23], and close to zero across the whole sample, *r* = 0.07, *t*(48) = 0.511, *p* = 0.611, 95% CI [-0.21; 0.35]. However, the difference in correlation between both groups was not significant, Δ*r*_*C<I*_ = 0.44, *z* = 1.520, *p* = 0.129, 95% CI [-0.13; 0.94].

Overall, these results suggest that there is an interaction between the incentives manipulation and aHPC-VTA/SN-RSFC change in how they relate to behavioural measures of learning, i.e., the total number of items encoded. To examine the robustness, Spearman’s rank correlation was used instead of Pearson’s correlation coefficient to calculate brain-behaviour correlations. Additionally, behavioural measures of learning were quantified in alternative ways (see Supporting Information and Figure S1 therein). For both behavioural indices of learning, the same results were observed when using Spearman’s rank correlation or alternative behavioural indices. We further examined whether the incentives effects on brain-behaviour correlations using the total number of items encoded as a behavioural measure of learning were observed independent of smoothing kernels (see Supporting Information and Table S1 and Figure S2 therein). The results showed that the effect was robust regardless of the smoothing kernels. Note that, when an FWHM kernel of 6 or 8 mm was used, Pearson’s correlation coefficients within the incentives group reached significance.

To determine whether the effect reported here was uniquely linked to aHPC-VTA/SN-RSFC change related to post-encoding connectivity, we controlled for aHPC-VTA/SN-FC during encoding (i.e., when participants watched magic tricks in the scanner, not during the resting-state scans). Specifically, we used a linear regression predicting each behavioural variable of interest using group, aHPC-VTA/SN-RSFC change, and the interaction between them. In line with the results reported above, the interaction term was the only significant predictor of the total number of items encoded and group was the only significant predictor of curiosity-driven memory benefit (see Supporting Information and Table S2 for details).

### Exploratory Whole-Brain FC Analysis

To examine the specificity of our results, exploratory whole-brain seed-to-voxel FC analyses were run to identify clusters showing a significant effect of the incentives manipulation in their change from pre- to post-learning rest in FC with the aHPC seed and the VTA/SN seed, respectively. Additional analyses were run to determine clusters in which the incentives manipulation influenced how the behavioural covariates of interest predicted the change from pre- to post-learning rest in FC with either seed. However, no clusters survived thresholding in any exploratory analysis.

## Discussion

We were interested to determine whether the same dopaminergic consolidation processes previously described in the context of reward-motivated learning (Gruber et al., 2016; Poh et al., 2021) can also be identified using a curiosity-motivated learning paradigm. As such, we investigated whether (1) there was an overall increase in RSFC between VTA/SN and aHPC as a function of learning, (2) change in RSFC would differ depending on the availability of extrinsic incentives, and (3) individual differences therein would predict behavioural measures of learning (i.e., the total number of items encoded and the curiosity-driven memory benefit). We did not find evidence for an overall increase in RSFC or for an effect of the incentives manipulation therein. However, our results showed that change in RSFC predicted the total number of items encoded differently depending on the availability of monetary incentives.

### No Statistically Significant Overall Change in aHPC-VTA/SN-RSFC or Incentives Effects

We found that overall, there was no significant increase in aHPC-VTA/SN-RSFC change from pre- to post-learning rest, neither across the whole sample nor as a function of the availability of incentives. The lack of overall change in RSFC between the aHPC and VTA/SN is somewhat surprising given the general assumption that changes in interregional FC from pre- to post-learning rest are related to the encoding, due to the post-learning rest being contrasted against the pre-learning rest baseline (Tambini & Davachi, 2019). While the lack of significant changes could imply that the encoding did not alter patterns of FC between the two areas that could reflect patterns of replay during consolidation processes (Josselyn et al., 2015), the lack of statistically significant effects indicating an overall shift in RSFC with encoding experience aligns with previous studies investigating changes in RSFC between MTL and dopaminergic midbrain (Gruber et al., 2016; Tompary et al., 2015). The absence of any effects of the incentives manipulation upon changes in RSFC suggests that providing additional extrinsic incentives in and of itself does not impact post-encoding communication between aHPC and VTA/SN. This is therefore somehow contradicting electrophysiological recordings in rodents showing an increase in hippocampal ripples after rewarded trials (Singer & Frank, 2009). However, it is important to note that electrophysiological recordings of hippocampal sharp wave ripples and aHPC-VTA/SN-RSFC change measured using fMRI BOLD responses are different measures. For instance, in the absence of an overall aHPC-VTA/SN-RSFC change from pre- to post-learning, Gruber and colleagues (2016) found an increase in hippocampal reactivation activity as a function of encoding, suggesting that overall local activation patterns can change without necessarily impacting systems-level interactions.

### Brain-Behaviour-Correlations are Affected by the Availability of Extrinsic Incentives

Our main finding is that the availability of extrinsic monetary incentives significantly impacted how intersubject variability in RSFC changes is associated with one of the behavioural measures of learning. More specifically, the correlation between the total number of items encoded and RSFC change between aHPC and VTA/SN differed significantly between both groups with larger brain-behaviour-correlations in the incentivised group (numerically positive correlation coefficient) compared to the control group (numerically negative correlation coefficient). The effects of curiosity on memory, on the other hand, were quantified using the curiosity-driven memory benefit. While the curiosity-driven memory benefit was significantly larger in the control compared to the incentivised group, no effects of the incentives manipulation on the brain-behaviour-correlations were found using the curiosity-motivated memory benefit as index. The findings were robust across different computation methods for correlation coefficients and various smoothing kernels. These effects were also robustly observed even after accounting for FC between aHPC and VTA/SN during online and offline encoding. As such, this suggests that the association between aHPC and VTA/SN FC in early consolidation during quiet awake rest posits a unique mechanism supporting learning rather than a continuation of FC processes observed during encoding. This observation is thereby in line with others deriving similar conclusions about the unique contribution of systems-level interaction during early consolidation to memory formation (Gruber et al., 2016; Murty et al., 2017; Tompary et al., 2015).

Overall, our results add to previous findings showing that individual differences in systems-level interactions differentially or selectively predict high-reward memory, in comparison to low-reward memory, where these effects were interpreted in light of a selective stabilisation of high-value memories (Cohen et al., 2021; Gruber et al., 2016; Murty et al., 2017). Other studies (Cohen et al., 2021; Gruber et al., 2016; Murty et al., 2017) investigating consolidation processes supporting memories for high-value information have manipulated value (i.e., incentives for intentional encoding or rewards in an unrelated task during incidental encoding) using within-subject manipulations. On the other hand, our study used a between-subject manipulation where half of the participants were informed that correct responses would be associated with additional monetary bonus payments. The adopted between-subject design has two implications. First, while a within-subject design has the advantage of higher statistical power, the design provides only one resting-state measure per participant, and it is difficult to tell whether the change reflects the encoding process of low- or high-value items. In contrast, the between-subject design allows us to obtain RSFC measures separately for the incentives and control conditions.

Second, in a within-subject design, it is likely that the cue indicating high- or low-value trials elicits a phasic dopamine response, i.e., a burst of activity in dopaminergic neurons. Such bursts facilitate the encoding of the dopamine-releasing events by cellular mechanisms of learning through the synthesis of plasticity-related protein during the expression of late LTP, which are then captured by tagged synapses, thereby enhancing their strength. These processes are then reflected in enhanced memory at the behavioural level (Düzel et al., 2010; Shohamy & Adcock, 2010). Importantly, similar phasic dopamine responses have been surmised to underlie curiosity (Bromberg-Martin & Hikosaka, 2009; Gruber et al., 2014; Gruber & Ranganath, 2019). Comparatively, in our study, these effects are assumed to be related to dopamine, but more particularly, to sustained changes in dopamine release, also known as tonic dopamine responses (Shohamy & Adcock, 2010). Such tonic dopaminergic responses are also assumed to be related to the effects of novelty exposure on encoding: Viewing novel scenes for 5 min has prolonged enhancement effects on the encoding of words studied afterwards (Fenker et al., 2008). However, rather than directly providing plasticity-related proteins to tagged synapses, prolonged tonic upregulation of dopamine might enhance the salience of environmental stimuli by making the dopaminergic neurons more likely to engage in phasic responses, increasing the likelihood that a presented stimulus will lead to a phasic response that then triggers plasticity-dependent protein synthesis processes (Düzel et al., 2010). Likewise, tonic dopaminergic activity might also influence hippocampal encoding processes within its own merits (Shohamy & Adcock, 2010). Overall, it is likely that the effects of dopamine on encoding cannot fully be attributed to phasic or tonic dopaminergic activity and are likely to be relying on a combination of these processes (Düzel et al., 2010; Shohamy & Adcock, 2010). In any case, if the incentives manipulation is indeed associated with tonic upregulation of dopaminergic activity as has been observed in the context of novelty, this might be the underlying mechanism that leads to the group effects observed and reported here.

While we found that brain-behaviour-correlations were significantly different as a function of the availability of extrinsic incentives, we did not find any evidence that curiosity-driven memory is related to aHPC-VTA/SN RSFC. Curiosity is often described as a motivation to reduce novelty and uncertainty or sometimes even as a motivation ‘for its own sake’ (Silvia, 2012). As such, stimuli that elicit curiosity can have a motivational salience within their own merit that can be similar to that found in the context of extrinsic incentives and rewards (FitzGibbon et al., 2020; Lau et al., 2020). Given previous studies showing that aHPC-VTA/SN-RSFC change supports memory for motivationally salient events in the context of extrinsic rewards (Cohen et al., 2021; Gruber et al., 2016), and proposals that extrinsic rewards and curiosity share features of incentive salience (FitzGibbon et al., 2020; Lau et al., 2020), it is tempting to predict that curiosity- and reward-motivated learning would use similar neural post-encoding mechanisms. However, we did not find that aHPC-VTA/SN-RSFC change predicted curiosity-driven memory benefit, neither across the whole sample nor within each group separately. This may simply reflect the issue of statistical power (sensitivity analysis indicated that the whole sample in this study was sufficient to detect moderate effects of *r* = 0.37 at the statistical power of 80%). Alternatively, the results may suggest that curiosity-motivated learning might be relying on, at least partially, post-encoding mechanisms other than dopaminergic influences on hippocampal activity measured using aHPC-VTA/SN-RSFC change. This is in line with behavioural results suggesting that the facilitating effects of curiosity on encoding occur independently of delay (Stare et al., 2018).

There are a few limitations to note. First, we aimed to investigate dopaminergic consolidation processes in the context of curiosity-motivated learning and how this is influenced by the availability of extrinsic incentives. Our analysis hence focused on the interaction between aHPC and VTA/SN because these regions are key players in models of motivated learning and have been implicated in the context of curiosity and extrinsic rewards/incentives (Gruber & Ranganath, 2019; Lisman et al., 2011; Lisman & Grace, 2005; Murty & Dickerson, 2016; Shohamy & Adcock, 2010). Effects within and between HPC and VTA/SN are often seen as an indication of dopaminergic activity. Indeed, it has been shown that dopamine release in the VTA/SN is reflected in the BOLD response (D’Ardenne et al., 2008). However, the VTA/SN also contains GABAergic and glutamatergic neurons (Nair-Roberts et al., 2008), which could be a source of the BOLD response (Düzel et al., 2009). Concluding with certainty that observed changes in the BOLD responses and FC are dopaminergic is hence impossible. Overall, the approach taken here is correlational in nature. It points towards the co-existence of learning and changes in RSFC, but it does not allow causal inferences regarding the role of dopamine. For this purpose, other methods are needed, including neurostimulation or pharmacological interventions. While pharmacological studies have been conducted in the context of reward and learning, most prominently in the context of instrumental learning (for a review, see Webber et al., 2021), pharmacological studies in the context of curiosity-motivated learning are thus far lacking and more research is needed in this area.

Second, in a related manner, studies targeting the dopaminergic midbrain and HPC also suffer from an overall poorer signal-to-noise ratio in these compared to cortical areas, making it even more challenging to capture effects in these areas. Indeed, as shown in the MMC Dataset Data Descriptor (Meliss, Pascua, et al., 2022), the temporal signal-to-noise ratio is lower in MTL and midbrain compared to e.g., the visual cortex. Likewise, prior literature has found evidence for reactivation within the entorhinal cortex (Staresina et al., 2013), systems-level consolidation between HPC and perirhinal cortex interacting with striatal activity (Murty et al., 2019), but also memory-reward interactions during encoding in the parahippocampal gyrus (Wolosin et al., 2012). Overall, this suggests that in addition to the HPC, other MTL structures might be involved in consolidation processes in the context of motivated learning. High-resolution fMRI studies (e.g., Duncan et al., 2014; Tompary et al., 2015; Wolosin et al., 2012) could help further disentangle the contributions of regions within the MTL, not only during consolidation but also during encoding, to enhance our understanding of motivated learning. More fine-grained approaches could also help to identify unique mechanisms in the context of curiosity- and extrinsically-motivated learning.

Third, the sample size of the study needs to be taken into consideration. While the sample size used here reflects those of typical fMRI studies (Szucs & Ioannidis, 2020), the sample size does not allow for reliable effect estimates in the context of brain-wide association studies, i.e., studies mapping individual differences in (f)MRI measures to cognitive function (Marek et al., 2022). In fact, while we found some near-significant results, we cannot know whether these results reflect the null results or the mere lack of statistical power. Hence, the robustness of the results needs to be confirmed in future studies.

In conclusion, the lack of a statistically significant association between aHPC-VTA/SN-RSFC change and curiosity-motivated learning potentially suggests that curiosity- and extrinsically-motivated learning do not rely on the same mechanisms during consolidation and that their effects on learning might thereby be additive. Indeed, the availability of extrinsic incentives influenced brain-behaviour correlations, suggesting that the availability of extrinsic incentives is associated with a large-scale switch in neural processes. In the absence of behavioural effects, however, we were unable to determine whether this switch is adaptive or maladaptive in nature and more research is needed to inform educational policies.

## Notes

The authors declare that they have no conflicts of interest in the subject matter or materials discussed in this manuscript.

We are grateful to the Centre for Integrative Neuroscience and Neurodynamics (CINN) and the MeMo Lab for helpful discussions and support in the realisation of this project. We would like to especially thank Cristina Pascua Martin for her support during the MRI data collection. We would like to express our gratitude to AFNI Team for their input regarding the fMRI pre-processing and analysis. Lastly, we would like to acknowledge Anthony Haffey’s ongoing support with regards to data collection using Collector. This research was supported by Leverhulme Trust Research Leadership Award (RL-2016-030), Jacobs Foundation Advanced Research Fellowship, and the Alexander von Humboldt Foundation (the Alexander von Humboldt Professorship endowed by the German Federal Ministry of Education and Research) to Kou Murayama.

https://doi.org/10.18112/openneuro.ds004182.v1.0.0

https://neurovault.org/collections/13320/

https://github.com/stefaniemeliss/MIC_paper

### Competing Interest Statement

The authors have declared no competing interest.

