## Supporting Information for "Curiosity-Motivated Incidental Learning With And Without Incentives: Early Consolidation And Midbrain-Hippocampal Resting-State Functional Connectivity"

### Alternative Behavioural Indices

In alignment with previous studies (Duncan et al., 2014; Tompary et al., 2015), we also computed a corrected memory score for each subject [high-confidence correct - (high-confidence incorrect)/4] to account for a larger proportion of incorrect response options in the four-alternative forced choice recognition memory test. Uncorrected and corrected number of items encoded were highly correlated, $r = 0.986$, $t(48) = 40.49$, $p < 0.001$. The use of an uncorrected or corrected total number of items encoded did not impact the results (see Figure S1 (left)).

We also examined whether an alternative way to quantify the effects of curiosity on memory would produce similar results. Specifically, we computed the within-person point-biserial correlation between high-confidence recognition performance (dummy-coded) and curiosity (ratings centred within clusters, i.e., each participant) across all 36 trials and Fisher's z-transformed the correlation coefficient before correlating them with aHPC-VTA/SN-RSFC change. Curiosity-driven memory benefit and within-person correlation coefficients were highly correlated, $r = 0.853$, $t(48) = 11.3$, $p < 0.001$ and the use of either of them did not impact the results reported (see Figure S1 (right)).

### Effects of Smoothing

In fMRI pre-processing, smoothing is usually carried out to increase the temporal signal-to-noise ratio (tSNR), however, with resting-state data, separating signal from noise remains challenging, making smoothing in the context of FC analysis a topic of debate. Investigating the impact of smoothing on RSFC, Molloy and colleagues (2014) did not find a significant effect of smoothing comparing an FWHM of 5.5mm to unsmoothed data, while Wu and colleagues (2011) reported that RSFC increased with increasing smoothing kernels with a significant jump between FWHM kernels of 4 and 6 mm. Recently, in a comparison between no smoothing and FWHM kernels of 4 and 8 mm, it has been shown that ROI-to-ROI-RSFC is not affected by the size of the kernel whereas effects were observed in seed-to-voxel analyses (Alahmadi, 2021). In all cases, conventional smoothing algorithms were applied. Critically, however, it has been pointed out that smoothing might be needed when the normalisation to template space involves non-linear transformations as this might introduce additional spatial effects and the usage of *'3dBlurToFWHM'* together with a small kernel to target small brain areas or a slightly larger kernel than the original imaging resolution to target cortical networks has been recommended (Wu et al., 2011). Importantly, increased head motion is correlated with intrinsic smoothness of the data, but smoothing using *'3dBlurToFWHM'* and an FWHM kernel of 6mm has also been shown to reduce motion confounds in RSFC (Scheinost et al., 2014). Due to conflicting results and recommendations to analyse data with different smoothing kernels to confirm the robustness of obtained results (Alahmadi, 2021), we here use *'3dBlurToFWHM'* with three different FWHM kernels (4, 6, and 8) as well as unsmoothed data to investigate the generalizability of results.

Results showed that overall aHPC-VTA/SN-FC estimates numerically increased as the smoothing kernel increased and that smoothing increased the variability within the FC estimates as well as for the change in RSFC. However, the overall results (e.g., whether a significant RSFC change or group effects therein were observed) were not affected. Likewise, when looking at brain-behaviour correlations, the results showed robust patterns (see Figure S2 and Table S1 below). This suggests that while the chosen smoothing kernel might influence the FC estimates, the individual differences therein seem to be preserved.

**Online and Offline Encoding**

Previous research showed that FC between HPC and VTA/SN during encoding is related to memory performance (Adcock et al., 2006; Duncan et al., 2014; Gruber et al., 2014; Shohamy & Wagner, 2008). However, these studies have used static stimuli like pictures or trivia questions. In the context of dynamic stimuli like video clips, however, a distinction can be made between online (intrastimulus) encoding processes, i.e., processes during stimulus presentation, and offline (poststimulus) encoding processes, i.e., processes during the period immediately following the event offset (Ben-Yakov & Dudai, 2011). In the majority of studies in cognitive neuroscience investigating the neural underpinnings of encoding, both processes are overlapping due to the use of discrete stimuli; however, the processes are more distinct in the context of dynamic stimuli. Indeed, it has been suggested that offline activity in the HPC following the offset of video clips can predict their encoding, either because the post-stimulus activity represents activity that binds the events of the clip into a single episode or because it reflects early consolidation (Ben-Yakov et al., 2013, 2014; Ben-Yakov & Dudai, 2011).

To determine whether any effects in changes in RSFC between aHPC and VTA/SN and behavioural measurements of encoding were unique to the early consolidation phase rather than reflecting general patterns during online and offline encoding, we also computed FC between aHPC and VTA/SN for each subject using the concatenated time series representing online and offline encoding, respectively, as input. fMRI timeseries acquired during the motivated incidental learning phase was used to compute FC between aHPC and VTA/SN during online and offline encoding.

In the motivated incidental learning phase, the beginning of magic tricks and fixations following them were aligned with the beginning of a TR. The duration of any fixations and response windows were multiples of the TR. While the fixation after the magic trick was jittered, the minimum duration was set to 4s (or 2 TRs). Because magic tricks were presented in pseudo-randomised order, the task time series was concatenated after pre-processing to remove volumes of no interest (e.g., ratings) and to reorder the volumes so that the final concatenaed time series would be the same across subjects (see Thomas et al., 2018) while accounting for the delay in the HRF by shifting the time course by a lag of 4 TRs (for details, see Meliss et al., 2022). This was done to create two separate time series: the online encoding time series covered only volumes acquired during the presentation of magic tricks (without the mock video) whereas the offline encoding time series included the last volume of each magic trick as well as the first two volumes of fixation after the end of the magic trick. The online time series consisted of 594 volumes and the offline time series consisted of 108 volumes with 36 volumes overlapping in both time series. Akin to the resting-state data, GCOR was computed for the online and offline encoding timeseries.

For each denoised time series, FC between aHPC and VTA/SN was determined akin to what has been described in the context of pre- and post-learning rest phases (i.e., by using *'3dSetupGroupInCorr'* and *'3dGroupInCorr'* accounting for the GCOR measurement obtained for each time series). As such, the FC measurements obtained for the online and offline encoding time series are similar to what others described as background FC[1] (Duncan et al., 2014; Murty et al., 2017, 2019; Tompary et al., 2015) which has been shown to reveal processes beyond typical task-based functional connectivity measures (Duncan et al., 2014) like beta-series correlation (Rissman et al., 2004). Importantly, however, no difference was calculated. Online and offline FC between aHPC and VTA/SN were significantly correlated, $r = 0.47$, $t(48) = 3.673$, $p = 0.006$, 95% CI [0.22; 0.66]. Lastly, all FC measures (change in RSFC, online FC and offline FC), as well as the availability of extrinsic incentives and the interaction thereof with change in RSFC were used to predict each behavioural measure of learning to determine the robustness of effects. As shown in Table S2, after accounting for all other variables, the interaction term between group and aHPC-VTA/SN-RSFC change remained the only significant predictor of the total number of items encoded. With respect to curiosity-driven memory benefit and in line with the results reported above, the availability of extrinsic incentives was the only significant predictor and no effects were observed for the interaction term between group and aHPC-VTA/SN-RSFC change.

[1] Studies using background functional connectivity typically apply event-related fMRI with simple, static stimuli. To remove stimulus- or response-induced neural activity, trial-evoked activity is removed using a Generalised Linear Model. This step was omitted due to the added complexity of modelling stimulus onset and duration in the context of naturalistic stimuli like video clips.

**Supplementary Tables**

Table S1

*Robust Correlation Coefficients Between aHPC-VTA/SN-RSFC Change and Behavioural Measures of Learning Irrespective of Pre-processing*

| | Whole sample | | Control group | | Incentives group | | Difference C < I | |
|---|---|---|---|---|---|---|---|---|
| | *r* | *p* | *r* | *p* | *r* | *p* | *r* | *p* |
| | | | | Total # items encoded | | | | |
| FWHM = 0 | 0.069 | 0.633 | -0.335 | 0.102 | 0.388 | 0.055 | 0.723 | 0.012 |
| FWHM = 4 | 0.08 | 0.58 | -0.313 | 0.128 | 0.389 | 0.054 | 0.702 | 0.015 |
| FWHM = 6 | 0.134 | 0.353 | -0.342 | 0.094 | 0.492 | 0.013 | 0.834 | 0.003 |
| FWHM = 8 | 0.165 | 0.253 | -0.328 | 0.11 | 0.536 | 0.006 | 0.864 | 0.002 |
| | | | | Curiosity-driven memory benefit | | | | |
| FWHM = 0 | 0.058 | 0.69 | -0.166 | 0.427 | 0.22 | 0.29 | 0.386 | 0.194 |
| FWHM = 4 | 0.074 | 0.611 | -0.18 | 0.389 | 0.269 | 0.193 | 0.449 | 0.129 |
| FWHM = 6 | 0.113 | 0.435 | -0.108 | 0.606 | 0.273 | 0.186 | 0.382 | 0.197 |
| FWHM = 8 | 0.122 | 0.399 | -0.092 | 0.662 | 0.279 | 0.177 | 0.371 | 0.209 |

*Note*. Correlation coefficients were computed using data from the whole sample as well as for each group individually and the difference in correlation between both groups. The same analysis was applied to data from different pre-processing pipelines using different FWHM kernels where 0 is equivalent to unsmoothed data and the other numbers represent the input for AFNI's 3dBlurToFWHM algorithm. *r* = Pearson correlation coefficient. *p* = p value.

Table S2

*Results of Linear Regression Predicting Behavioural Measures of Learning*

| | Whole sample | | Control group | | Incentives group | |
|---|---|---|---|---|---|---|
| | *b* (SE) | *p* value | *b* (SE) | *p* value | *b* (SE) | *p* value |
| | | | Total # items encoded | | | |
| Intercept | 15.75 (1.06) | < 0.001 | 15.52 (0.94) | < 0.001 | 14.76 (1.16) | < 0.001 |
| Online FC | -20.1 (41.51) | 0.631 | 15.36 (48.47) | 0.755 | -74.39 (74.79) | 0.331 |
| Offline FC | 16.59 (29.85) | 0.581 | 0.15 (37.75) | 0.997 | 30.12 (47.37) | 0.532 |
| Incentives | -0.77 (1.5) | 0.611 | | | | |
| RSFC change | -49.44 (34.18) | 0.155 | -45.68 (31.55) | 0.162 | 84.79 (42.11) | 0.057 |
| Incentives * RSFC change | 127.24 (49.12) | 0.013 | | | | |
| | | | Curiosity-driven memory benefit | | | |
| Intercept | 0.1 (0.04) | 0.028 | 0.08 (0.04) | 0.057 | -0.05 (0.04) | 0.284 |
| Online FC | -3.01 (1.65) | 0.075 | -1.7 (2.13) | 0.435 | -4 (2.64) | 0.145 |
| Offline FC | 0.82 (1.19) | 0.494 | -1.15 (1.66) | 0.496 | 2.86 (1.67) | 0.102 |
| Incentives | -0.14 (0.06) | 0.023 | | | | |
| RSFC change | -1.32 (1.36) | 0.336 | -0.81 (1.39) | 0.567 | 1.99 (1.49) | 0.196 |
| Incentives * RSFC change | 3.63 (1.96) | 0.07 | | | | |

*Note.* The table above shows the results of the linear regressions predicting each behavioural measure of learning using the availability of extrinsic incentives ("Incentives", effect-coded using 1 for incentives and -1 for control group), aHPC-VTA/SN-RSFC change ("RSFC change"), and their interaction ("Group * RSFC change") after controlling for the FC between aHPC and VTA/SN during encoding ("Online FC" and "Offline FC") across the whole sample. Given the significant interaction term for both measures, a linear regression was run within each group using Online and Offline FC as well as RSFC change as predictors to further understand the interaction effects. *b* = unstandardised beta coefficient. SE = standard error.

**Supplementary Figures**

Figure S1

*Brain-Behaviour Correlations Using  Alternative Ways to Quantify Behavioural Measures of Learning*

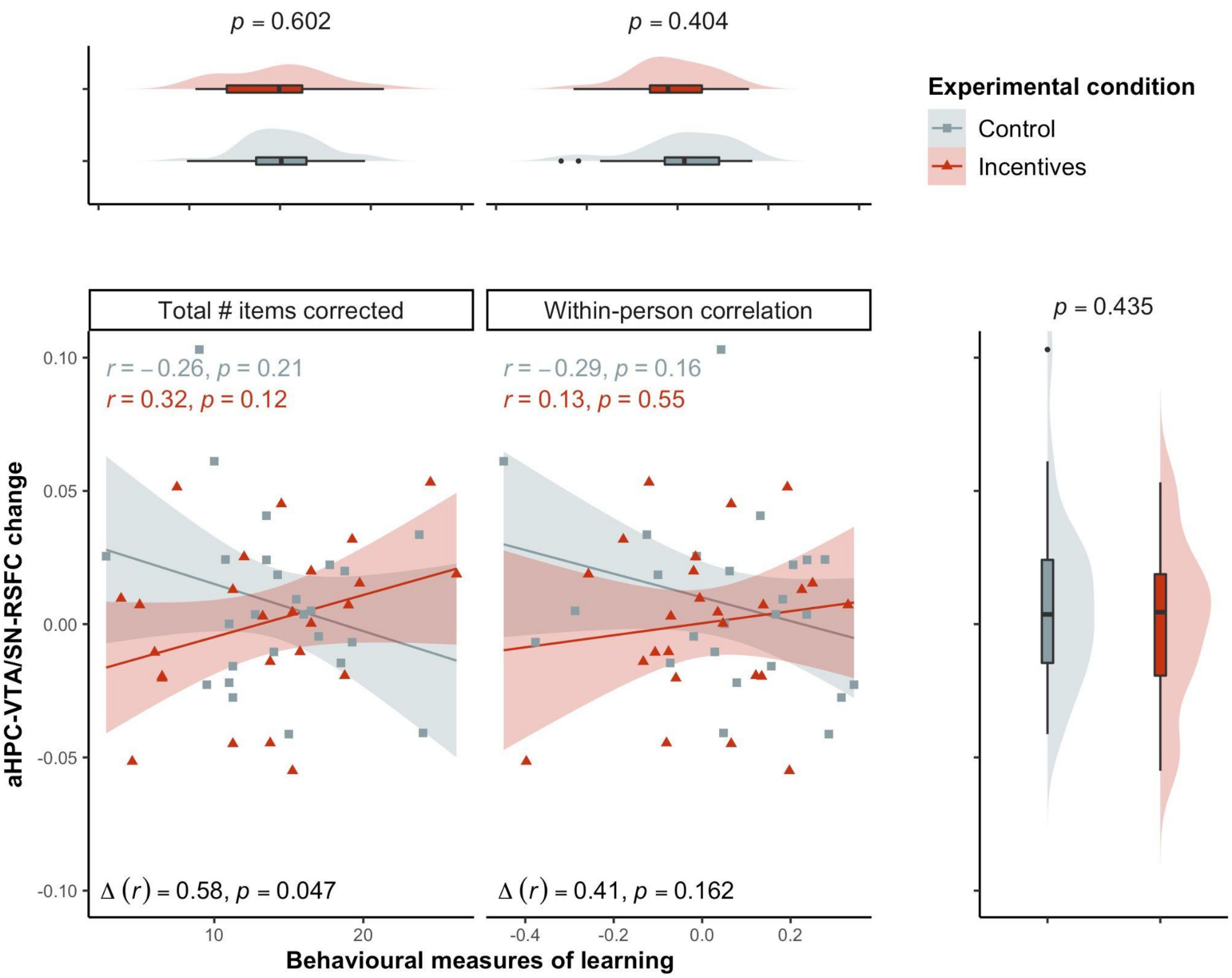


*Note*. The scatter plots above show the association between behavioural measures of learning on the x-axis and changes in FC between aHPC and VTA/SN from pre- to post-learning rest on the y-axis. Data is shown separately for the corrected total number of items encoded (left) and the within-person correlations between curiosity and memory (right). Different colours were used for the control and the incentives group, respectively. Regression lines with 95% confidence have been added to illustrate the relationship between the behavioural measures of learning and aHPC-VTA/SN-RSFC change within each group together with the corresponding correlation coefficients.

Figure S2

*Brain-Behaviour Correlation Coefficients as a Function of Learning Phase and Pre-processing*

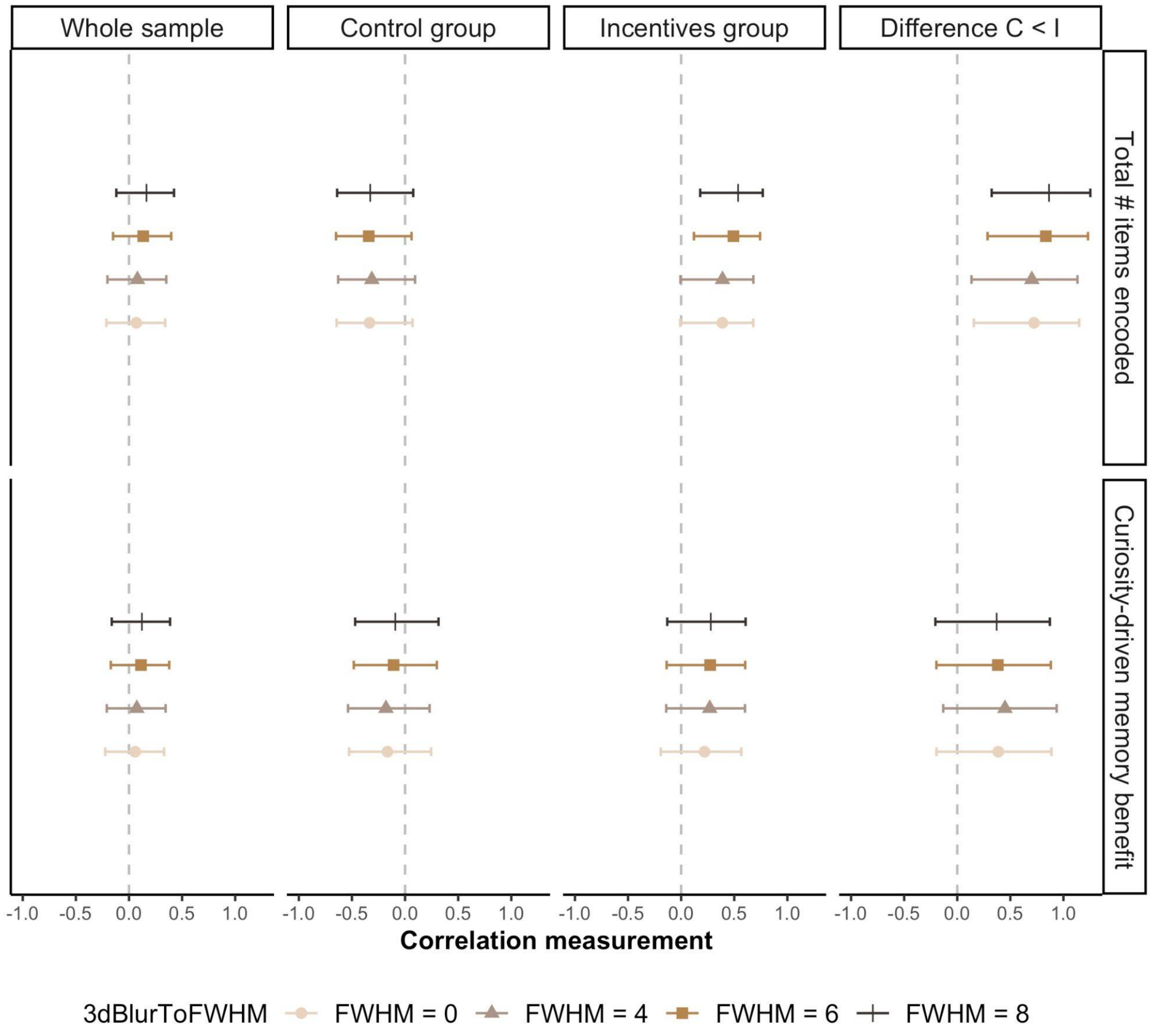


*Note*. The plot shows the computed correlation between behavioural measures of learning and aHPC-VTA/SN-FC measures obtained during different phases of the task. Correlation coefficients are computed using data from the whole sample as well as for each group individually and the difference in correlation between both groups. Error bars represent the 95% CI. As behavioural measures of learning the total number of items encoded (top) and curiosity-driven memory benefit (bottom) were used. FC measures between aHPC and VTA/SN were computed separately for each pre- and post-learning rest phase to determine the change therein. Different colours denote FHWM kernels used in the pre-processing where 0 is equivalent to unsmoothed data and the other numbers represent the input for AFNI's *3dBlurToFWHM* algorithm.
